# A deep learning-based multisite neuroimage harmonization framework established with traveling-subject dataset

**DOI:** 10.1101/2021.12.05.471192

**Authors:** Dezheng Tian, Zilong Zeng, Xiaoyi Sun, Qiqi Tong, Huanjie Li, Hongjian He, Jiahong Gao, Yong He, Mingrui Xia

## Abstract

The accumulation of multisite large-sample MRI datasets collected by large brain research projects in the last decade has provided a critical resource for understanding the neurobiological mechanisms underlying cognitive functions and brain disorders. However, the significant site effects, observed in the imaging data and their derived structural and functional features, has prevented the derivation of consistent findings across different studies. The development of harmonization methods that can effectively eliminate complex site effects while maintaining biological characteristics in neuroimaging data has become a vital and urgent requirement for multisite imaging studies. Here, we proposed a deep learning-based framework to harmonize imaging data from pairs of sites, in which site factors and brain features can be disentangled and encoded. We trained the proposed framework with a publicly available traveling-subject dataset from SRPBS and harmonized the gray matter volume maps from eight source sites to a target site. The proposed framework significantly eliminated inter-site differences in gray matter volume. The embedded encoders successfully captured both the abstract texture of site factors and the concrete brain features. Moreover, the proposed framework exhibited outstanding performance relative to conventional statistical harmonization methods in site effect removal, data distribution homogenization, and intra-subject similarity improvement. Together, the proposed method offers a powerful and interpretable deep learning-based harmonization framework for multisite neuroimaging data that could enhance reliability and reproducibility in multisite studies for brain development and brain disorders.

## 1 Introduction

Advances in magnetic resonance imaging (MRI) in the recent decades have offered potent techniques for noninvasively exploring the structures and functions of the human brain *in vivo*, leveraging our understanding for the neurobiological mechanisms underlying the development of complex cognitions and the clinical impairments related to brain disorders (Cao et al., 2017a; Fornito et al., 2015; Park and Friston, 2013). The practice of multisite MRI data acquisition in recently launched large brain research projects, such as the IMAGEN (Schumann et al., 2010) and ABCD (Casey et al., 2018), has accumulated critical neuroimaging resources to facilitate brain investigation with impressive statistical power (Laird, 2021; Poldrack and Gorgolewski, 2014; Xia and He, 2017). However, considerable heterogeneity among imaging datasets collected from different sites, that is, the site effect, has been widely documented, in both the raw structural and functional imaging data (Li et al., 2020; Radua et al., 2020) and image-derived brain characteristics, such as gray matter volume (GMV) (Melzer et al., 2020) and functional connectivity (Noble et al., 2017a; Yamashita et al., 2019). The site effect predominantly results from both the sampling of divergent populations and the different scan equipment across different sites and is a major source of the inconsistencies in the findings reported from different studies on the same topic. Therefore, developing methods for harmonizing imaging data across different scan sites has become a fundamental and urgent requirement for multisite imaging studies.

To correct for the site effect in multisite imaging data, several harmonization strategies have been proposed, which can be summarized into two major categories: conventional statistics-based harmonization methods and recently developed deep learning (DL)-based harmonization methods. Conventional statistical methods are usually applied in a linear regression manner on univariate metrics with sites indexed as a categorical covariate, for example, the least squares-based general linear model (Rao et al., 2017) and Bayesian estimation-based ComBat (Fortin et al., 2018; Fortin et al., 2017). These methods have been utilized in multisite imaging studies and have shown a powerful capacity for removing linear site effects in brain metrics (Pomponio et al., 2020; Xia et al., 2019; Yu et al., 2018). However, noticeable limitations have been observed for this type of harmonization method. First, the site effect is mathematically assumed to be linear, while the actual effect may be much more complex. Second, brain characteristics are considered independently in these models, largely neglecting the spatial and topological relationships among brain regions. To overcome these defects, recently proposed DL-based harmonization methods, including U-net (Dewey et al., 2019), cycle-generative adversarial network (Modanwal et al., 2020), or three-dimensional convolutional neural network (Tong et al., 2020), allow for mapping the complex abstract representations of the nonlinear spatial pattern of the site effects in MRI data. These models have been primarily applied to the harmonization of diffusion tensor images (Moyer et al., 2020), structural images (Zuo et al., 2021), and morphological measurements (Zhao et al., 2019), successfully eliminating the site effect in such data with complex spatial or topological information. However, the interpretability is relatively low for most of these established DL-based harmonization methods, for which high-dimensional representations are difficult to delineate. Additionally, the model training strategy of site pairing is a common approach for DL-based methods, and the fusion of data from multiple sites in a single model will greatly increase the model’s complexity and require much more training data. Designing a harmonization framework with high expandability will facilitate the application of DL-based methods.

Another critical factor for establishing reliable multisite image harmonization models is the selection of training data. The core objectives of multisite harmonization are the elimination of non-biological factors, such as MRI equipment and scan protocols, while simultaneously retaining the biological factors of the participants across different sites. Therefore, the innovative traveling-subject dataset, in which each participant is scanned at all different sites, has become a valuable resource for the training of harmonization models, as it can minimize the bias of population sampling across sites and ensure that the established models only learn non-biological factors (Noble et al., 2017b; Tong et al., 2019; Yamashita et al., 2019). Although existing multisite imaging studies have shown that harmonization models based on nontraveling-subject datasets, be they conventional statistics or DL-based models, can efficiently remove the site effect (Garcia-Dias et al., 2020), it is unknown whether the inter-site differences in biological factors are over eliminated. Benefiting from the publicly available traveling-subject dataset, several recent studies have established harmonization methods that can separate and protect biological factors from complex site effects and have achieved outstanding performance with a small training sample (Yamashita et al., 2019). However, DL-based harmonization models for brain measurements using traveling-subject dataset are still lacking.

Here, we proposed a DL-based harmonization framework that can disentangle both site-factor and brain-factor representations from site effects based on a publicly available traveling-subject dataset. Taking the widely used GMV measurement as an illustration, we first examined whether this framework can significantly eliminate site effects in the GMV maps of nine scan sites. Then, we investigated whether the site-factor and brain-factor encoders embedded in the framework can capture inter-site and inter-subject variability, respectively. Finally, we compared proposed methods with several conventional statistical harmonization methods in terms of site effect removal, data distribution homogenization, and intra-subject similarity improvement.

## 2 Methods

### 2.1 The deep learning-based representation disentanglement (DeRed) framework for multisite imaging data harmonization

We proposed a DL-based bidirectional framework (Fig. 1a) for neuroimaging data harmonization, which enables the transfer of imaging data from a given site to a target site, and vice versa. Specifically, this framework contains four encoders for disentangling site-factors and brain-factors in imaging data of the source and target sites, and two decoders for synthesizing the harmonized images for the encoders. This design allows harmonized imaging data to contain both target site information and natural brain features. This framework was inspired by a disentangled unsupervised cycle-consistent adversarial network (DUNCAN) (Liu et al., 2021), which was developed to remove MRI artefacts based on representation disentanglement. As shown in Fig. 2a, the site-factor encoder in DeRed is designed to have three residual blocks which can avoid the convergence performance degradation caused by structure redundancy (He et al., 2016a, b). Each residual block includes a set of 2D-Convonlution Layer, and leaky rectified linear unit (LeakyReLU) activation (Fig. 2b). When the feature maps pass through the residual block, the size is reduced by half, and the output of each residual block can be used as image features at different scales. Notably, each input slice of the site-factor encoder must undergo an average pooling process before the residual blocks because the representation related to the scanning site or equipment should be abstract, regardless of anatomical details, and should not be extracted from the shallower layer. Similar to the site-factor encoder, the brain-factor encoder is composed of four residual blocks. The difference is that the brain-factor encoder lacks the average pooling process and each residual block contains the instance normalization operation (Huang and Belongie, 2017) after LeakyReLU activation to capture independent features across imaging data of the same subject.

**Fig. 1.**
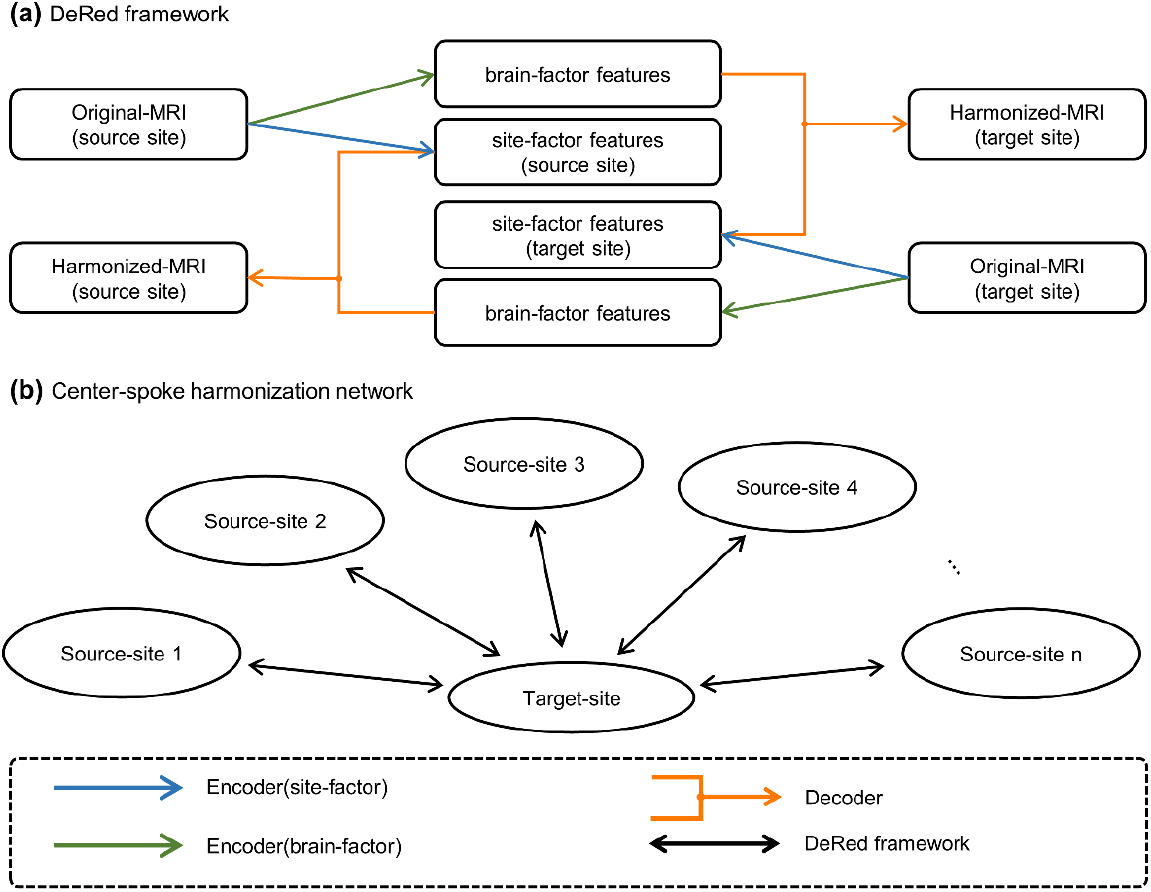
Architecture of the DeRed framework and center-spoke harmonization network. **(a)** The DL-based representation disentanglement framework. The site-factor and brain-factor features are extracted from original MR images by the encoders, and the decoder synthesizes harmonized MR images by combining these two features. **(b)** Center-spoke harmonization network with the target site located at the center. This harmonization network supports the bidirectional migration of MRI between the target site and the source sites.

**Fig. 2.**
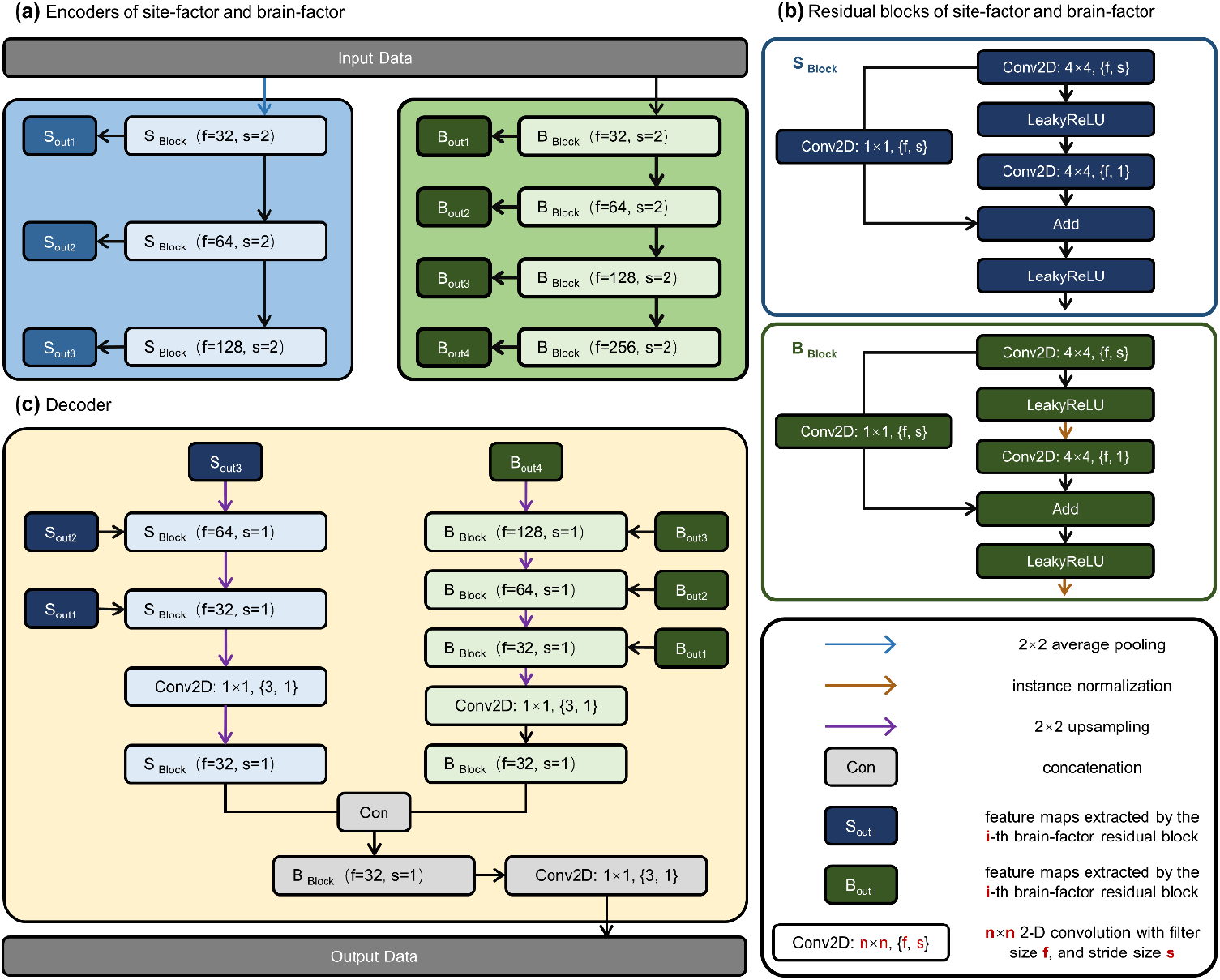
Architecture of the encoders and decoder. **(a)** Architecture of the site-factor encoder and brain-factor encoder. The S_out i_ and B_out i_ models represent the feature maps extracted by the *i*-*th* site-factor and brain-factor residual block, respectively. **(b)** Architect of the site-factor residual block (S _Block_) and brain-factor residual block (B _Block_). **(c)** Architecture of the decoder, which integrating the outputs from both site-factor and brain-factor encoders.

The decoder (Fig. 2c) contains a two-step synthesis structure, integrating features extracted by the encoders. First, the site-factor features at different scales are mixed through a series of upsampling processes and residual blocks, it should be noted that the size of each feature map is not be reduced by half when passing through the residual block. Similarly, the mixing process for brain-factor features also involves brain-factor residual blocks and an upsampling process. After the first stage of the mixing process, the decoder produces two feature maps, one for the site-factors and the other for the brain-factors. Second, the mean, maximum and minimum feature maps are calculated, and these feature maps are concatenated and input into a brain-factor residual block with a 2D-convonlution operation. The input data of the site-factor encoder and the brain-factor encoder are two-dimensional images obtained by slicing along a certain direction of within three-dimensional data, and the output results of the decoder maintain a consistent shape with the input data.

Based on the DeRed framework, we established a flexible harmonization network as shown in Fig. 1b. The different sites can be understood as different nodes in this harmonization network, connected by edges played by DeRed. The harmonization network possesses a center-spoke topology, with the target site, whose scanned images have the best data quality, at the center, and the data of th other sites are harmonized to this center site. Notably, the scanning data from any site can be transferred to another site through the network edges. Furthermore, if a new site establishes a relationship with a site belonging to this harmonization network, it can also be transferred to any other site along the network edges.

### 2.2 Materials and T1 data processing

To minimize sampling bias across sites, we trained our harmonization framework using a traveling-subject dataset from the DecNef Project Brain Data Repository (https://bicr-resource.atr.jp/srpbsts/), which was gathered by the Japanese Strategic Research Program for the Promotion of Brain Science (SRPBS) (Tanaka et al., 2021; Yamashita et al., 2019). This dataset included nine healthy participants (all male, ages 24-32 years), each of whom underwent T1-weighted MRI scans at 12 different centers. All of these sites used 3T scanners but with different manufacturers (Siemens, GE, and Philips), scanner types (Verio, Tim Trio, Spectra, Skyra, and Achieva), phase encoding directions (posterior to anterior and anterior to posterior), and numbers of channels per coil (8, 12, and 32). Data from three sites were excluded (ATT, UTO, YC2) due to duplicate data. The detailed scanning parameters at each site are listed in Table S1.

In the current study, we selected the widely used GMV measurement (Grieve et al., 2013; Smallwood et al., 2013) derived from T1-weighted images as an example to examine the feasibility of the proposed harmonization method. The calculation of the GMV was carried out by using Statistical Parametric Mapping (SPM12, https://www.fil.ion.ucl.ac.uk/spm/) (Ashburner, 2012) and the Computational Anatomy Toolbox (CAT12, http://dbm.neuro.uni-jena.de/cat12/) (Iglesias et al., 2015). Briefly, for each T1 scan, an N4 bias field inhomogeneity correction was first performed, and an adaptive maximum a posteriori (AMAP) approach was then used in tissue segmentation. Optimized shooting approach-based spatial registration was further performed to normalize all images into the standard Montreal Neurological Institute (MNI) space. Modulated normalization was then implemented to compensate for GMV changes caused by affine transformation and nonlinear warping. Finally, all GMV maps were smoothed with an 8-mm full-width at half-maximum (FWHM) Gaussian kernel.

### 2.3 Training and harmonization processes

ATV was selected as the target site (*φ_t_*) in the harmonization process mainly for the following two reasons. First, the equipment manufacturer and number of channels per coil of ATV were the most frequently used among all the sites. Second, the imaging data from ATV showed better quality with less noise than those from other sites according to the visual screening. Other sites were regarded as the source sites (*φ_s_*), resulting in 8 independent inter-site harmonization pairs with ATV.

Prior to the training process, we cropped all the GMV maps from a matrix size of (181, 217, 181) to (176, 208, 176), which guaranteed that the sliced images could can be restored to their original size after multiple average pooling and upsampling operations. Moreover, to ensure the harmonization process within the gray matter regions and reduce the computational burden, we constrained the data training process within a gray matter mask, which was determined by averaging the GMV maps of all scans and further applying a threshold of 0.2 mm^3^.

The inputs of the training model were obtained by slicing along a certain anatomical direction (coronal, sagittal, or transverse); slices that did not intersect with the gray matter mask were not included in the subsequent training process. A section position was then randomly determined during each epoch to ensure uncertainty during the training process, and slices of imaging data of all subjects at *φ_t_* and *φ_s_* were extracted at this position. Notably, we hold that spatially adjacent slices assist in capturing brain-factor representation information, so we set the spatial resolution of the training slices to (176, 208, 3) for the transverse orientation, (176, 176, 3) for the coronal orientation, and (208, 176, 3) for the sagittal orientation. Thus, the *i-th* individual slice can be predicted repetitively at different channels for the (*i-1*)-*th*, *i-th* and (*i*+*1*)-*th* slice inputs. The images resulted from the three channels were averaged to obtain the final harmonized single slice.

Furthermore, if the harmonization process is simply based on a single slicing direction, it cannot fully summarize the global spatial information of 3D-images. Therefore, we independently trained three models. The training set of each model was obtained by slicing the image data from different anatomical directions, and then the output was averaged as the final harmonization result of the 3D image.

We defined four convergence constraint losses for the harmonization procedure:

First, we expect the site-factor encoder to extract the same representation at the same site across different subjects:

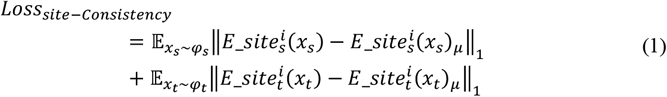

where *x_s_* and *x_t_* denote the images from *φ_s_* and *φ_t_*, respectively. 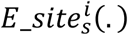 and 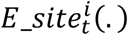 denote the *i*-*th* feature map outputs of the *i*-*th* residual block in the site-factor encoders of *φ_s_* and *φ_t_*, respectively. 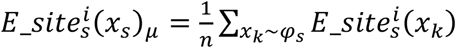 and 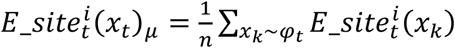 denote the average *i-th* site-factor residual block outputs of n subjects from *φ_s_* and *φ_t_*, respectively.

Second, we expect the brain-factor encoders of *φ_s_* and *φ_t_* to extract the same representation from imaging data acquired from the same person but at different sites:

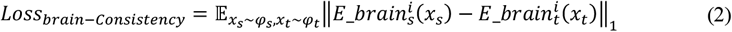

where 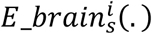 and 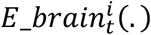 denote the *i*-*th* feature map outputs of the *i*-*th* residual block in the brain-factor encoders from *φ_s_* and *φ_t_*, respectively.

Third, we encourage the decoders to reconstruct the images by merging the site-factor representation and the brain-factor representation from their own sites. This self-reconstruction loss can be formulated as:

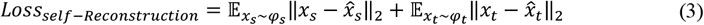

Fourth, the site-factor representation from *φ_t_* is necessary for the decoder in *φ_t_* to reconstruct the images, even if the brain-factor representation belongs to *φ_s_*. In the same way, the decoder of *φ_s_*can reconstruct images according to the site-factor representation from *φ_s_* and the brain-factor representation from *φ_t_*. The cross-reconstruction loss can be formulated as

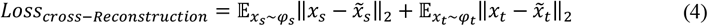

where 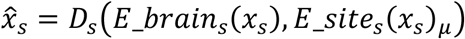(*E_brain_s_*(*x_s_*), *E_site_s_*(*x_s_*)*μ*) and 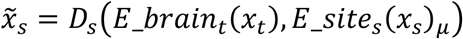 denote the reconstructed images, both of which contain the site-factor representation from *φ_s_* but the brainfactor representation from *φ_s_* and *φ_t_*, respectively. In contrast, 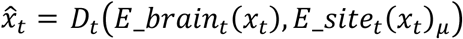 and 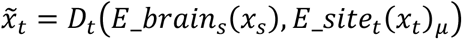 denote the reconstructed images, both of which contain site-factor representation from *φ_t_* but the brain-factor representation from *φ_t_* and *φ_s_*, respectively.

### 2.4 Evaluation of harmonization outcome

We trained the DeRed harmonization network with a total of 81 images from all subjects scanning across all sites and obtained the corresponding harmonization results, which were used to quantify the inter-site differences and explain the representation captured by DeRed.

#### 2.4.1 Correction for site effects

We adopted two methods to examine whether the proposed framework can reduce the site effects on the GMV maps. First, we performed linear discriminant analysis (LDA), a classic dimensionality reduction technique, to project the GMV measurement into two coordinates with the scanning site as a prior classification label. LDA is commonly used to project features into a lower dimension space by maximizing the distance between classes and minimizing the variation within each class. In this study, the site effect was reflected by the clustering of data from the same site. Second, we used one-way analysis of variance (ANOVA) to quantitatively test for significant site differences in the GMV. The significance level of the voxel-wise comparison was set to a voxel-level *p* < 0.001 with a cluster-level Gaussian random field (GRF)-corrected *p* < 0.05.

#### 2.4.2 Interpretability of the encoders

To assess whether each kind of encoder (i.e., site-factor and brain-factor) captured the corresponding features, we examined the output images by blocking their opposite input of the decoder in turn. To interpret the site-factor encoder, we set all values of the brain-factor feature maps to zero, and feed them into the decoder. The image synthesized in this case can be understood to contain only the site-factor representation (i.e., *I_site_*). Assuming that each site-factor encoder captures the characteristics of the scanner, the inter-site variance of *I_site_* should be spatially similar to the inter-site variance in the original GMV images. Thus, we first calculated the variance of each voxel of *I_site_* and averaged the GMV variance maps of each subject across sites. We then applied Spearman’s correlation to examine the spatial correlation of these two variance maps.

To interpret the brain-factor encoder, we fed the decoder with brain-factor feature maps and empty site-factor feature maps (i.e., feature maps with 0 values), and the image synthesized in this case can be understood to contain only the brain factor representation (i.e., *I_bram_*). To examine whether the brain-factor encoder truly captures these individual heterogeneity-related representations, we first assessed Spearman’s correlation for each voxel between the original GMV and age across subjects, preserving those voxel groups *S_α_* whose GMV was significantly correlated with age. Then, we also preserved those voxel groups *S_β_* with a significant correlation between *I_brain_* and age. The overlap between *S_α_* and *S_β_* was then calculated by 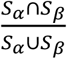. Second, the similarity between the original GMV and *I_brain_* was calculated for each subject according to the Spearman’s correlation coefficient.

### 2.5 Comparison between DeRed and other harmonization methods

Several harmonization methods have been proposed to remove site effect differences in recent multicenter studies, including general linear model harmonization (GLM), global scaling harmonization (GS), and ComBat harmonization (see SI for a detailed description of these methods). To examine the advantages of our proposed methods, we compared DeRed with these harmonization methods in terms of site effect removal, GMV distribution coherence, intra-subject similarity improvement and inter-subject difference reservation. A leave-one-subject-out cross-validation strategy was utilized for each method. Briefly, we excluded the data of the *i*-*th* subject at all sites, trained the framework with the remaining 72 scanned images from the other subjects, and applied the trained model to harmonize the data from the *i*-*th* subject. This procedure was repeated nine times to select each subject as the test data in turn.

#### 2.5.1 Site effect removal

To test whether site effects could be removed by all the methods, we used ANOVA on the harmonized GMV maps for each method. The significance level of the voxel-wise comparison was set to a voxel-level *p* < 0.001 with a cluster-level GRF-corrected *p* < 0.05. Furthermore, we used Wilcoxon signed-rank tests to compare the *F* values between the original and harmonized data, and between harmonization results from different methods.

#### 2.5.2 GMV distribution consistency

For the original data and harmonized data of each method, we first calculated the average GMV map across subjects and estimated their probability distribution for each site. We then estimated the averaged bidirectional KL divergence between each pair of probability distributions for different sites. The KL divergence was further compared between the original and harmonized data and between harmonization results from different methods with Wilcoxon signed-rank tests.

#### 2.5.3 Inter-subject difference reservation

The difference across subjects was calculated using the Euclidean distance of the original GMV maps within each site and further averaged across all sites to obtain a reference inter-subject difference matrix. Then, for each harmonization results from different methods, we calculated the inter-subject difference matrix within each site. Spearman’s correlation was further used to estimate the correlation between each matrix and the reference matrix. A significant correlation coefficient indicated the preservation of inter-subject differences.

#### 2.5.4 Intra-subject similarity improvement

For each subject, we calculated the Spearman’s correlation coefficient between the GMV map of any pair of sites among the nine sites as the intra-subject similarity. These correlation coefficients were then compared using Wilcoxon signed-rank tests between the original and harmonized data and between harmonization results from different methods.

## 3 Results

### 3.1 Site effect removal of DeRed

We first visualized the heterogeneity in the original and harmonized GMV maps across nine sites by projecting their dominant features into a 2D space using LDA decomposition. The site-clustered distribution of the LDA features indicated noticeable inter-site heterogeneity for the original GMV maps (Fig. 3a). Specifically, data from HUH and HKH were the most distant from other datasets, which might essentially be due to their unique scanner models (GE Signa HDxt for HUH and Siemens Sepctra for HKH). However, the harmonized data showed a relatively homogeneous distribution, implying the effective removal of the site effect (Fig. 3b). Subsequent statistical analysis confirmed this finding that one-way ANOVA revealed a significant site effect across the nine sites on the original GMV maps, primarily in the medial temporal and occipital cortices, the insula, and the cerebellum (Fig. 3c, voxel-level *p*<0.001, GRF-corrected *p*<0.05). In contrast, no significant site effect was observed in the harmonized GMV maps derived from our proposed DeRed framework (Fig. 3d, voxel-level *p*<0.001, GRF-corrected *p*<0.05). To further illustrate the order of scan properties (e.g., MRI manufacturer, scanner type, and phase coding) that contribute to the site effect, we performed a hierarchical clustering on regions showing significant site effect across nine sites. We found that the manufacturer of the scanner was the most distinguish factor for the site effect (Fig. S1).

**Fig. 3.**
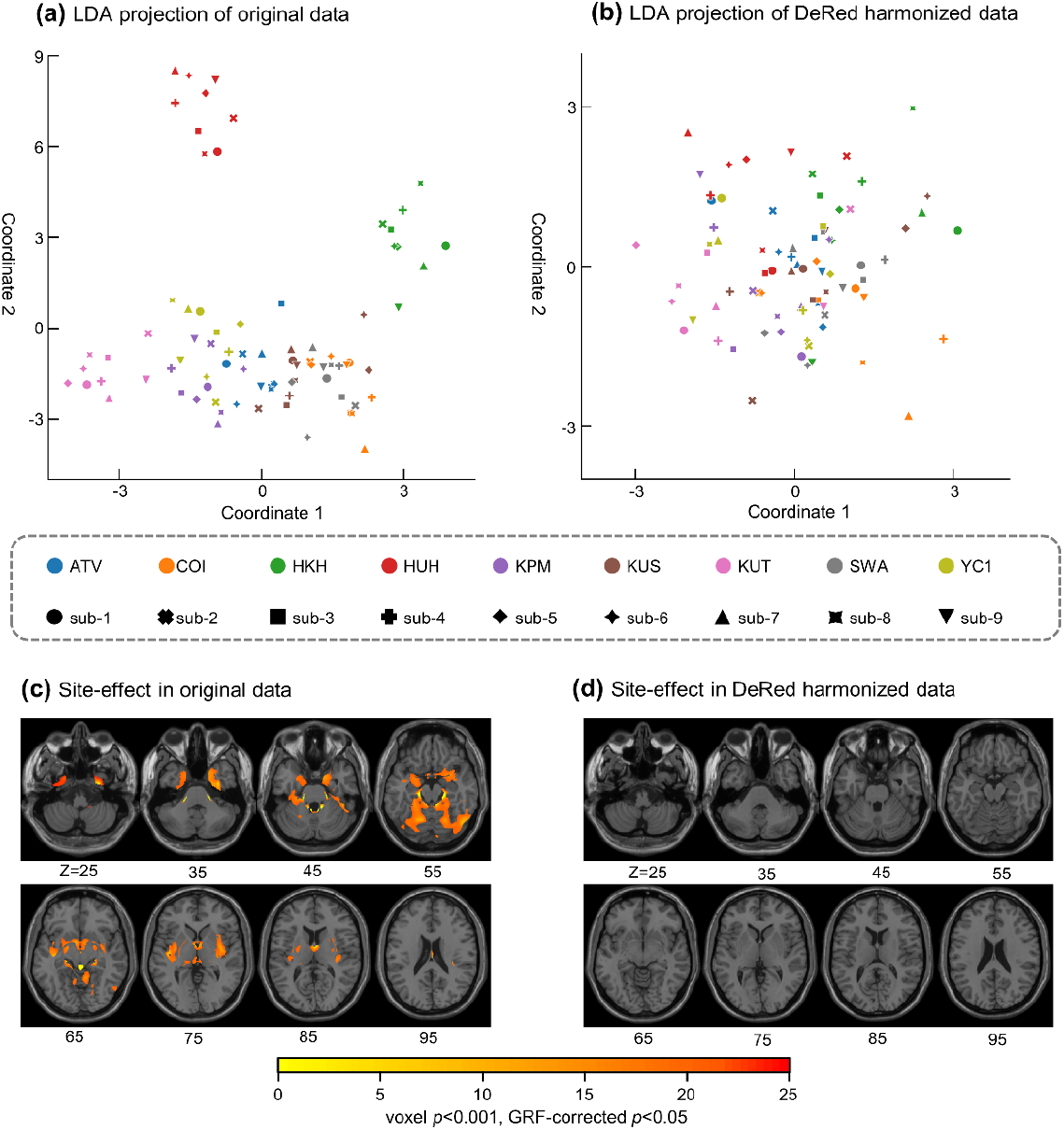
Site effect in data before and after harmonization. **(a)** and **(b)** illustrate LDA projection of GMV before and after harmonization. A datapoint represents a projected GMV measurement from a subject, its color represents the site from which it originates and its shape represents the subject to which it belongs. **(c)** and **(d)** illustrate the site effect identified by one-way ANOVA in the original and harmonized GMV respectively, sliced along the transverse anatomical orientation. There were no significant differences across sites for all voxels after DeRed harmonization.

### 3.2 Interpretability of the encoders

We examined the feature representation of the site-factor and brain-factor encoders by blocking their respective opposite outputs. As illustrated by randomly chosen data (e.g., sub-01 at YC1) in Fig. 4a, the outputs from the site-factor encoders were decoded into a field map with abstract boundaries of the brain and blurry texture on the background. In contrast, images decoded from the brain-factor encoders showed the detailed structure of the gray matter anatomy, which was highly similar to that of the original GMV maps. Further quantitative analysis showed that the inter-site variance of *I_site_* was significantly spatially correlated with the inter-site variance of the original images in the log-log coordinates (Fig. 4b, Spearman’s correlation, *ρ* =0.42, *p*<0.0001), suggesting that the site-factor encoder captures the variance of actual physical factors across the scanner. For the *I_brain_*, we first found that they were significantly spatially correlated with the original GMV maps for each individual in each site (Spearman’s correlation, *ρ* =0.993 ± 0.002, all *p*<0.0001). We then examined the overlap of clusters showing significant correlations with those in the original data. In the original data, we found that the GMV was significantly positively correlated with age in the right precuneus, inferior frontal gyrus, and the left parahippocampus, and negatively correlated with age mainly in the dorsolateral prefrontal, visual, and lateral temporal cortices (voxel-level *p*<0.001, GRF-corrected *p*<0.05). The *I_brain_* showed similar distributed brain-age correlations at all nine sites (Fig. 4c, overlap ratio of the significant voxels: 75.54% ± 2.43%). Together, these results suggest that the brain-factor encoders successfully captured the biological details of the individual GMV maps.

**Fig. 4.**
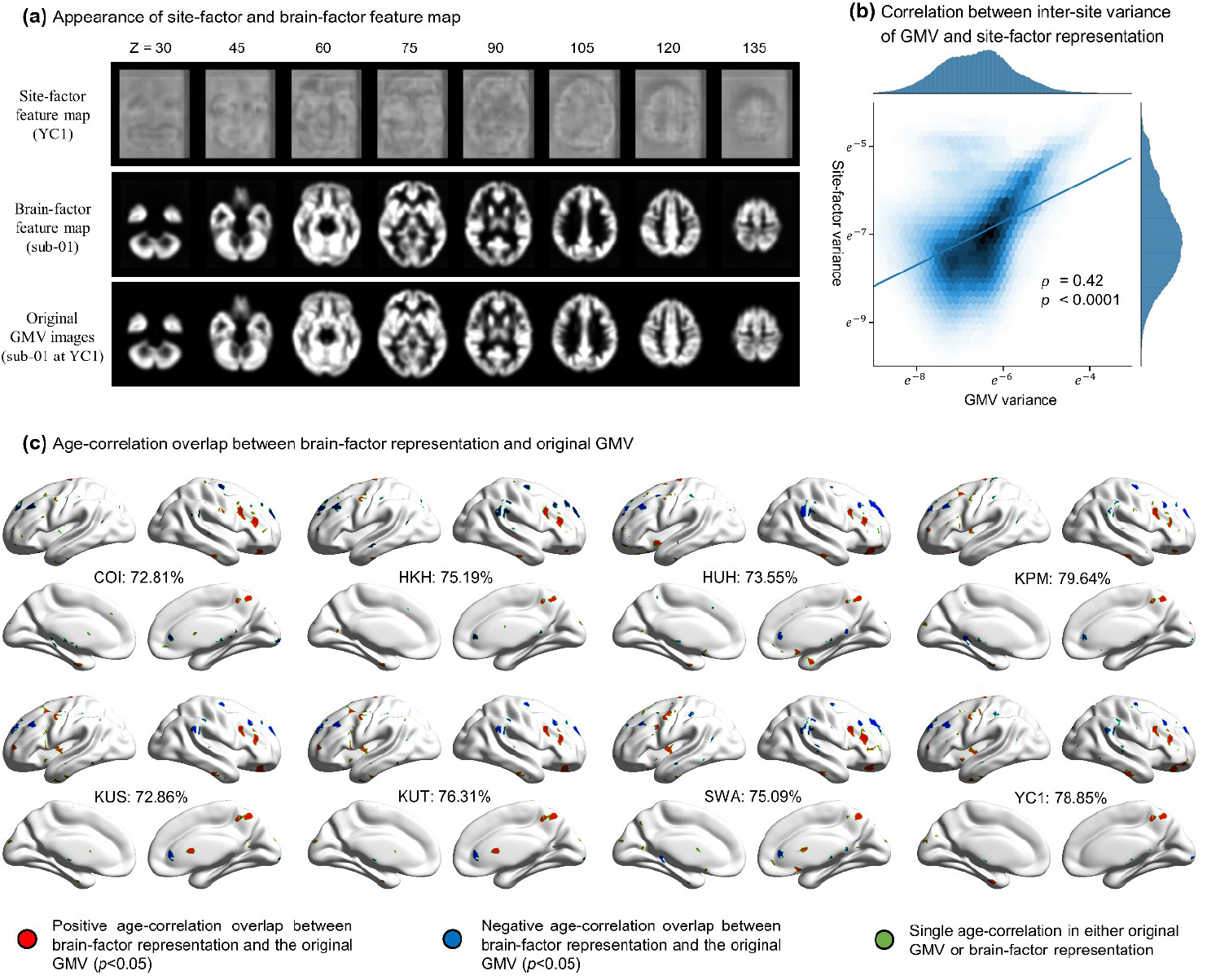
Interpretability of site-factor and brain-factor encoders. **(a)** Appearance of site-factor and brain-factor feature map. the first row represents the decoder outputs containing only site-factor representations of YC1. The second row represents the decoder outputs containing only brain-factor representations of subject-01. The last row represents the original GMV map of subject-01 from YC1. **(b)** Log-log correlation plot between the original GMV variance and the variance of the site-factor feature maps. Each variance measurement is transformed by natural logarithm conversion. The color depth reflects the dot density within a single hexagon. **(c)** Age-correlation overlap clusters between brain-factor representation and the original GMV. The voxels in red (blue) regions indicate a positive (negative) age correlation (*p*<0.05) in both the original GMV and the brain factor representation. These voxels colored green indicate a single age correlation in either the original GMV or brain-factor representation.

### 3.3 DeRed showed better harmonization performance than conventional methods

We compared the performance of the proposed DeRed harmonization framework to that of several conventional methods, including GS, GLM, and ComBat. First, we found that the significant site effects in the original data could be entirely eliminated by DeRed and ComBat but partly retained significant in the data processed with GS and GLM (Fig. 5a, ANOVA, voxel-level *p*<0.001, GRF-corrected *p*<0.05). Further between-method comparisons showed that the site effect (*F* value estimated in ANOVA) was significantly lower in the harmonized data from DeRed than in those from other methods (Fig. 5b, Wilcoxon signed-rank tests, *p* <0.001, Bonferroni-corrected).

**Fig. 5.**
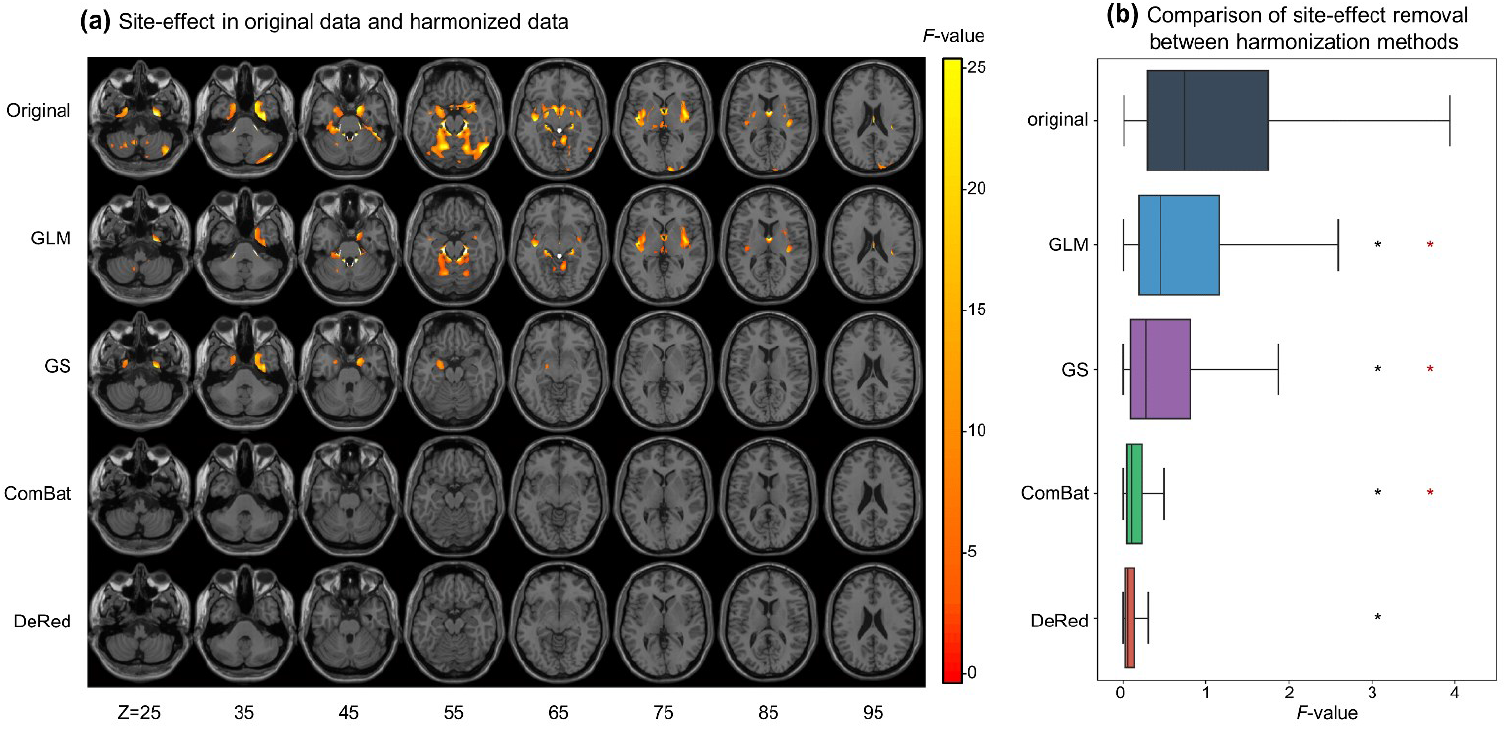
Site effect removal of different harmonization methods. **(a)** Site effect identified by ANOVA in data before and after harmonization (voxel-level *p*<0.001, GRF-corrected *p*<0.05). **(b)** Comparison of site effect (*F* value) in data before and after harmonization by different methods; all harmonization results show a lower *F* values than the original state (Wilcoxon signed-rank tests, *p*<0.05, Bonferroni-corrected, labeled by black asterisk), and the *F* values of DeRed harmonized data are significantly lower relative to those of other methods (Wilcoxon signed-rank tests, *p*<0.05, Bonferroni-corrected, labeled by red asterisk).

Second, we found that the probability distributions of the averaged GMV maps were divergent across the nine sites, and the distributions of the harmonized data tended to be more consistent (Fig. 6a).

**Fig. 6.**
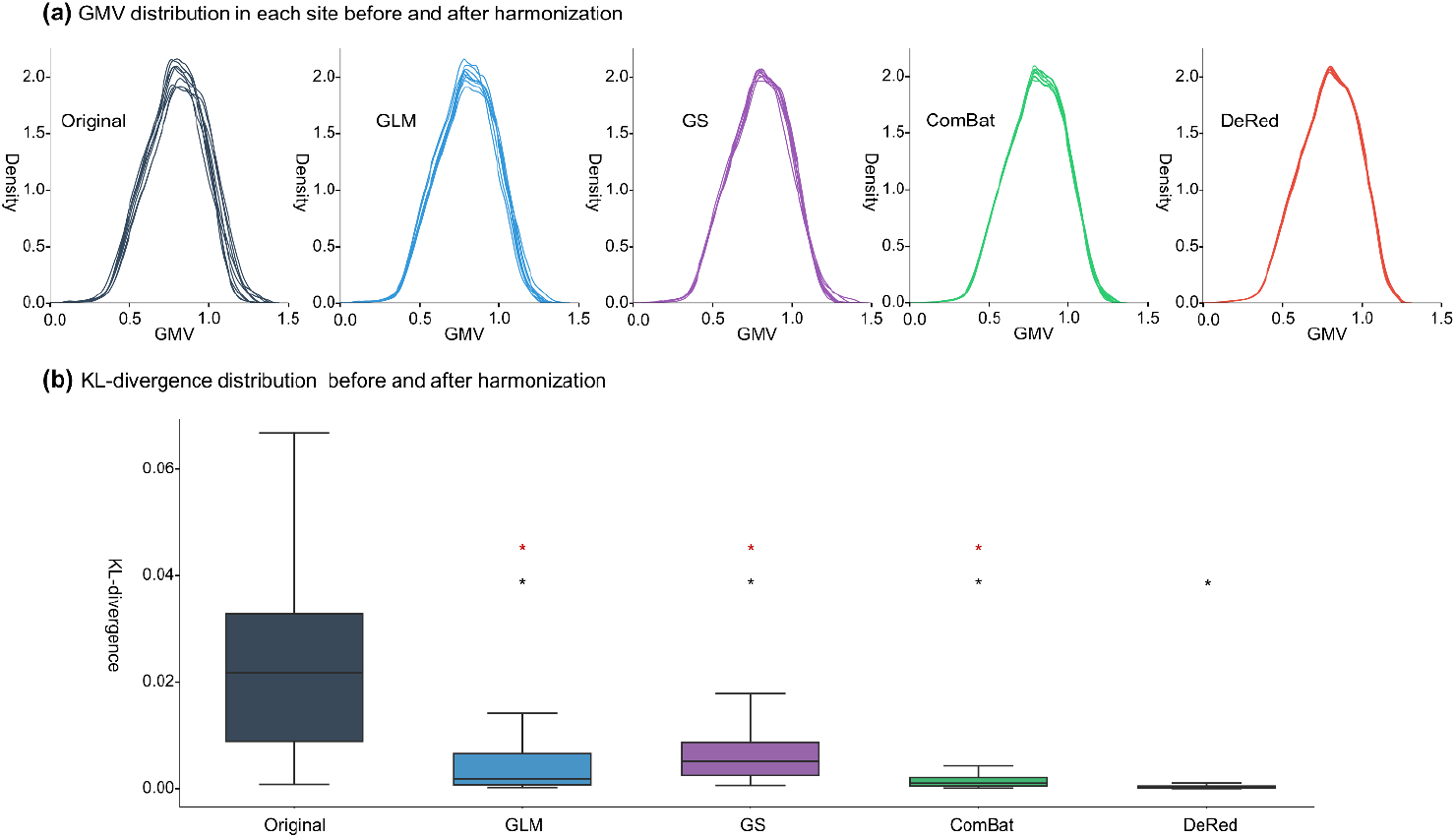
Divergence in the GMV distribution across different sites before and after harmonization. **(a)** GMV distribution in different sites. Each curve represents the probability distribution of the GMV measurement for all voxels averaged across subjects in a site. **(b)** Boxplots of KL-divergence across sites before and after harmonization by different methods. All harmonization data showed a lower JS-divergence compared with the original data (Wilcoxon signed-rank tests, *p*<0.05, Bonferroni-corrected, labeled by black asterisk). DeRed demonstrated a significantly lower KL-divergence relative to the compared methods (Wilcoxon signed-rank tests, *p*<0.05, Bonferroni-corrected, labeled by red asterisk).

Quantitatively, the harmonized data derived from all methods showed a significantly lower KL divergence than the original data, and data derived from DeRed exhibited the lowest KL divergence among the harmonization methods (Fig. 6b, Wilcoxon signed-rank tests, *p*<0.001, Bonferroni-corrected).

Finally, the inter-subject distance matrix for each site derived from the harmonized data for each method was significantly correlated with the original averaged matrix (Fig. 7a and Fig. S2, Spearman’s correlation, *ρ*=0.90 ± 0.04, all *p*<0.0001), indicating that all harmonization methods maintained the inter-subject differences in the GMV. Moreover, we found the intra-subject similarity on the GMV was significantly increased for all harmonization methods (Wilcoxon signed-rank tests, *p* <0.05, Bonferroni-corrected). Importantly, DeRed demonstrated the statistically highest intrasubject similarity among all harmonization methods (Fig. 7b, Wilcoxon signed rank tests, *p* <0.05, Bonferroni-corrected), indicating that the proposed framework has the greatest ability to increase intra-subject consistency across sites.

**Fig. 7.**
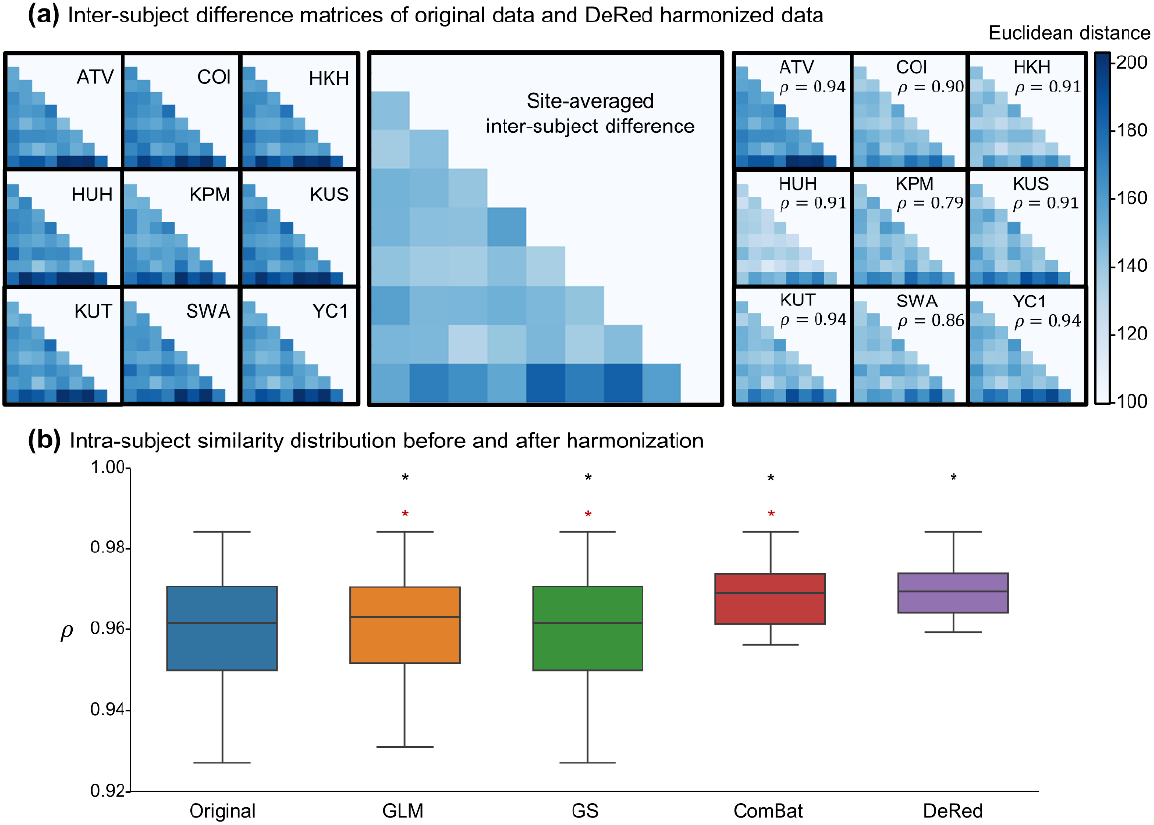
Inter-subject difference maintenance and intra-subject similarity improvement before and after harmonization. **(a)** Inter-subject difference matrix before and after harmonization at each site. The difference matrices were averaged across sites before harmonization. The color depth of *i*-*th* row and *j*-*th* column grid for each matrix represents the Euclidean distance between the *i*-*th* and *j*-*th* subjects. Spearman’s correlation coefficient is illustrated by *ρ* (*p*<0.001). (d) Boxplots of intra-subject similarity across sites before and after harmonization by different methods. The harmonization results with higher self-identifiability relative to the original data are labeled with black asterisks. DeRed demonstrated a significantly improved intra-subject similarity (*p*<0.05) over all comparison methods based on the paired Wilcoxon signed-rank test, with a red asterisk.

## 4 Discussion

In this paper, we proposed a DL-based harmonization framework for multisite MRI data named DeRed, which was further trained with a traveling-subject dataset. Taking the commonly used GMV metric as an example, the proposed framework showed good performance in eliminating the divergence in the GMV across different sites. Notably, the encoders embedded in the framework successfully captured both the abstract textures of site factors and the concrete biologically related brain features. Moreover, the proposed framework exhibited outstanding performance relative to conventional harmonization methods in site effect removal, data distribution homogenization, and intra-subject similarity improvement. Together, the proposed method offers a powerful and extendable DL-based harmonization framework for multisite neuroimaging data with high interpretability, facilitating the improvement of the reliability and reproducibility of multisite studies for brain development and brain disorders.

Compared with traditional statistics-based harmonization methods, the advantages of the proposed DL-based framework can be formulated from several perspectives. First, instead of taking a single metric as an independent variable, the DL model comprehensively extracts the global and local imaging information by integrating information from spatially neighboring units (e.g., voxels in a brain map) through a series of convolution and pooling operations (Bau et al., 2020). Many studies have suggested that adjacent voxels reflect closer correlations both in the anatomical structure and in the physiological mechanism (Cao et al., 2017b; Cigdem et al., 2019). These individual-specific anatomical details embodied within the MR images are repeatable across multisite measurements and should not be ignored during the harmonization process. Second, both DL-based methods and statistics-based methods attempt to explore the mapping relationship during the harmonization process. However, harmonization processes guided by statistical strategies, such as GLM and GS, seem to be limited in the ability to map linear polynomial functions. In our work, we employed the residual block inside the proposed framework, which has been shown to be especially important for fitting a more accurate function map mixed with a variety of high-dimensional and nonlinear characteristics between the MR images and the site effect representations (Lusch et al., 2018). Third, statistics-based harmonization frameworks scrupulously rely on the prior assumption. For example, ComBat describes the site effect of each voxel via additive and multiplicative factors, which are assumed to follow the normal distribution and inverse gamma distribution respectively (Johnson et al., 2007). Nevertheless, the site effect reflected within the MR images can be understood as a heterogeneous mixture caused by the action of an asymmetrical magnetic field and complex neurophysiological activity (Vovk et al., 2007), which is difficult to generalize adequately with simple probability distributions. Compared with statistics-based methods, the proposed harmonization framework driven by the pixel-to-pixel loss function, is not limited to the prior distribution assumption, allowing the harmonization results of DeRed to demonstrate a better probability consistency across sites.

In the current study, we mainly employed the voxel-based GMV, a commonly used structural brain measurement to validate our proposed framework. It should be noted that the proposed DL-based representation disentanglement and reconstruction strategy can be referenced to other multisite harmonization processes for structural and functional brain metrics in diverse formats. Regarding the volumetric images, our proposed framework can provide a robust contribution by fine tuning its network architecture. Similar designs to the encoders and the decoders embedded in DeRed were adopted to extract the latent representations and remove the artefacts in T1-w and T2-w MR images in a previous study (Liu et al., 2021). For those data in a network format, such as the structural and functional connectivity matrices, the graph convolutional network (GCN) can be integrated in the proposed framework. Many studies have applied the GCN to reveal functional brain network similarity, comprehensively considering its topological property (Ktena et al., 2018) and to more efficiently predict the longitudinal development of cognitive performances (e.g., motor and cognitive scores) of preterm infants by recognizing the local and global topology patterns of their structural brain network (Kawahara et al., 2017). Illuminated by existing studies, the GCN can be used to depict complex topological mechanisms and identify abstract high-dimensional information, indicating that the application of the GCN may help to capture the site-specific topological effect, from which multisite structural and functional brain network harmonization can be reasonably performed.

Several issues should be further considered. First, the proposed framework was trained on a traveling-subject dataset to minimize sampling bias across scan sites. However, the traveling-subject MRI data collection design is generally lacking in many multisite databases. Therefore, we intend to adopt random bootstrap sampling to produce a biological-matching dataset from each site (Kim et al., 2021) and further expand the proposed framework for unpaired inter-site datasets. Second, the traveling-subject dataset used in this work was acquired from a group of healthy participants aged from 24 to 32 years and the biological validation was limited due to the lack of cognitive or clinical evaluations; thus, the generalizability of DeRed to MRI data acquired from special populations (e.g., children and adolescents or patients with brain disorders) needs to be further validated. Studies have revealed significant development effects and disorder-related disruptions in brain structure and functions (Gilmore et al., 2018; van den Heuvel and Sporns, 2019). Therefore, the specific optimization strategy for harmonization methods needs further investigation for these special populations. Third, similar to most DL method, the proposed DeRed framework comprised several convolution and pooling operations. Therefore, the harmonized data could be objectively smoothed with neighboring information during encoding and decoding. Although this procedure overcomes local noise during harmonization, further validations for data distribution and design optimization for the DL network are required. Finally, in the current study, we preliminarily established a multisite harmonization network based on the DeRed framework, and its expandability needs further evaluation. Considering the bidirectional connectedness of this network, data can be harmonized to any node (e.g., site) in the network. Moreover, a new site could be easily included in the harmonization network by training a DeRed model between it and any existing sites.

**Table 1.** Details of the scanning parameters in the traveling-subject dataset.

| Site | ATV | COI | HKH | HUH | KPM | KUS | KUT | SWA | YC1 |
| --- | --- | --- | --- | --- | --- | --- | --- | --- | --- |
| Manufacturer | Siemens | Siemens | Siemens | GE | Philips | Siemens | Siemens | Siemens | Philips |
| Platform | Verio | Verio | Spectra | Signa HDxt | Achieva | Skyra | TimTrio | Verio | Achieva |
| Magnetic field strength (T) | 3.0 | 3.0 | 3.0 | 3.0 | 3.0 | 3.0 | 3.0 | 3.0 | 3.0 |
| Number of channels per coil | 12 | 12 | 12 | 8 | 8 | 32 | 32 | 12 | 8 |
| Phase encoding | PA | AP | PA | PA | AP | AP | PA | PA | AP |
| Echo time (ms) | 2.98 | 2.98 | 2.38 | 1.928 | 3.31 | 2.98 | 3.4 | 2.98 | 3.176 |
| Repetition time (ms) | 2300 | 2300 | 1900 | 6788 | 7.1 | 2300 | 2000 | 2300 | 6.99 |
| Flip angle (°) | 9 | 9 | 10 | 20 | 10 | 9 | 8 | 9 | 9 |
| Image dimension | 240×256×256 | 176×240×256 | 224×320×320 | 180×256×256 | 170×256×256 | 224×232×256 | 240×256×208 | 240×256×256 | 200×256×256 |
| Pixel dimension | 1×1×1 | 1×1×1 | 0.81×0.75×0.75 | 1×1×1 | 1×1×1 | 1×1×1 | 0.9375×0.9375×1 | 1×1×1 | 1×1×1 |

Abbreviations: ATV, Siemens Verio scanner at the Advanced Telecommunications Research Institute International; COI, Center of Innovation at Hiroshima University; HKH, Hiroshima Kajikawa Hospital; HUH, Hiroshima University Hospital; KPM, Kyoto Prefectural University of Medicine; KUS, Siemens Skyra scanner at Kyoto University; KUT, Siemens Tim Trio scanner at Kyoto University; SWA, Showa University; YC1, Yaesu Clinic scanner 1; PA, posterior to anterior; AP, anterior to posterior

## Acknowledgements

This work was supported by National Natural Science Foundation of China (Nos. 82071998, 82021004, 81671767, and 81620108016), Beijing Nova Program (No. Z191100001119023), Fundamental Research Funds for the Central Universities (No. 2020NTST29), the National Key R&D Program of China (No. 2018YFA0701400), and the Changjiang Scholar Professorship Award (No. T2015027).

## References

Ashburner, J., 2012. SPM: a history. Neuroimage 62, 791–800.

Bau, D., Zhu, J.Y., Strobelt, H., Lapedriza, A., Zhou, B., Torralba, A., 2020. Understanding the role of individual units in a deep neural network. Proc Natl Acad Sci U S A 117, 30071–30078.

Cao, M., Huang, H., He, Y., 2017a. Developmental Connectomics from Infancy through Early Childhood. Trends in Neurosciences 40, 494–506.

Cao, S., Li, Y., Deng, W. W., Qin, B.Y., Zhang, Y., Xie, P., Yuan, J., Yu, B.W., Yu, T., 2017b. Local Brain Activity Differences Between Herpes Zoster and Postherpetic Neuralgia Patients: A Resting-State Functional MRI Study. Pain Physician 20, E687–E699.

Casey, B.J., Cannonier, T., Conley, M.I., Cohen, A.O., Barch, D.M., Heitzeg, M.M., Soules, M.E., Teslovich, T., Dellarco, D.V., Garavan, H., Orr, C.A., Wager, T.D., Banich, M.T., Speer, N.K., Sutherland, M.T., Riedel, M.C., Dick, A.S., Bjork, J.M., Thomas, K.M., Chaarani, B., Mejia, M.H., Hagler, D.J., Cornejo, M.D., Sicat, C.S., Harms, M.P., Dosenbach, N.U.F., Rosenberg, M., Earl, E., Bartsch, H., Watts, R., Polimeni, J.R., Kuperman, J.M., Fair, D.A., Dale, A.M., Workgrp, A.I.A., 2018. The Adolescent Brain Cognitive Development (ABCD) study: Imaging acquisition across 21 sites. Developmental Cognitive Neuroscience 32, 43–54.

Cigdem, O., Demirel, H., Unay, D., 2019. The Performance of Local-Learning Based Clustering Feature Selection Method on the Diagnosis of Parkinson’s Disease Using Structural MRI. 2019 Ieee International Conference on Systems, Man and Cybernetics (Smc), 1286–1291.

Dewey, B.E., Zhao, C., Reinhold, J.C., Carass, A., Fitzgerald, K.C., Sotirchos, E.S., Saidha, S., Oh, J., Pham, D.L., Calabresi, P.A., van Zijl, P.C.M., Prince, J.L., 2019. DeepHarmony: A deep learning approach to contrast harmonization across scanner changes. Magnetic Resonance Imaging 64, 160–170.

Fornito, A., Zalesky, A., Breakspear, M., 2015. The connectomics of brain disorders. Nat Rev Neurosci 16, 159–172.

Fortin, J.P., Cullen, N., Sheline, Y.I., Taylor, W.D., Aselcioglu, I., Cook, P.A., Adams, P., Cooper, C., Fava, M., McGrath, P.J., McInnis, M., Phillips, M.L., Trivedi, M.H., Weissman, M.M., Shinohara, R.T., 2018. Harmonization of cortical thickness measurements across scanners and sites. Neuroimage 167, 104–120.

Fortin, J.P., Parker, D., Tunc, B., Watanabe, T., Elliott, M.A., Ruparel, K., Roalf, D.R., Satterthwaite, T.D., Gur, R.C., Gur, R.E., Schultz, R.T., Verma, R., Shinohara, R.T., 2017. Harmonization of multi-site diffusion tensor imaging data. Neuroimage 161, 149–170.

Garcia-Dias, R., Scarpazza, C., Baecker, L., Vieira, S., Pinaya, W.H.L., Corvin, A., Redolfi, A., Nelson, B., Crespo-Facorro, B., McDonald, C., Tordesillas-Gutierrez, D., Cannon, D., Mothersill, D., Hernaus, D., Morris, D., Setien-Suero, E., Donohoe, G., Frisoni, G., Tronchin, G., Sato, J., Marcelis, M., Kempton, M., van Haren, N.E.M., Gruber, O., McGorry, P., Amminger, P., McGuire, P., Gong, Q.Y., Kahn, R.S., Ayesa-Arriola, R., van Amelsvoort, T., de la Foz, V.O.G., Calhoun, V., Cahn, W., Mechelli, A., 2020. Neuroharmony: A new tool for harmonizing volumetric MRI data from unseen scanners. Neuroimage 220.

Gilmore, J.H., Knickmeyer, R.C., Gao, W., 2018. Imaging structural and functional brain development in early childhood. Nat Rev Neurosci 19, 123–137.

Grieve, S.M., Korgaonkar, M.S., Koslow, S.H., Gordon, E., Williams, L.M., 2013. Widespread reductions in gray matter volume in depression. Neuroimage Clin 3, 332–339.

He, K.M., Zhang, X.Y., Ren, S.Q., Sun, J., 2016a. Deep Residual Learning for Image Recognition. 2016 Ieee Conference on Computer Vision and Pattern Recognition (Cvpr), 770–778.

He, K.M., Zhang, X.Y., Ren, S.Q., Sun, J., 2016b. Identity Mappings in Deep Residual Networks. Computer Vision - Eccv 2016, Pt Iv 9908, 630–645.

Huang, X., Belongie, S., 2017. Arbitrary Style Transfer in Real-time with Adaptive Instance Normalization. 2017 Ieee International Conference on Computer Vision (Iccv), 1510–1519.

Iglesias, J.E., Augustinack, J.C., Nguyen, K., Player, C.M., Player, A., Wright, M., Roy, N., Frosch, M.P., McKee, A.C., Wald, L.L., Fischl, B., Van Leemput, K., Neuroimaging, A.D., 2015. A computational atlas of the hippocampal formation using ex vivo, ultra-high resolution MRI: Application to adaptive segmentation of in vivo MRI. Neuroimage 115, 117–137.

Johnson, W.E., Li, C., Rabinovic, A., 2007. Adjusting batch effects in microarray expression data using empirical Bayes methods. Biostatistics 8, 118–127.

Kawahara, J., Brown, C.J., Miller, S.P., Booth, B.G., Chau, V., Grunau, R.E., Zwicker, J.G., Hamarneh, G., 2017. BrainNetCNN: Convolutional neural networks for brain networks; towards predicting neurodevelopment. Neuroimage 146, 1038–1049.

Kim, N., Arfanakis, K., Leurgans, S.E., Yang, J.Y., Fleischman, D.A., Han, S.D., Aggarwal, N.T., Lamar, M., Yu, L., Poole, V.N., Bennett, D.A., Barnes, L.L., 2021. Bootstrap approach for meta-synthesis of MRI findings from multiple scanners. Journal of Neuroscience Methods 360.

Ktena, S.I., Parisot, S., Ferrante, E., Rajchl, M., Lee, M., Glocker, B., Rueckert, D., 2018. Metric learning with spectral graph convolutions on brain connectivity networks. Neuroimage 169, 431–442.

Laird, A.R., 2021. Large, open datasets for human connectomics research: Considerations for reproducible and responsible data use. Neuroimage 244.

Li, H.J., Smith, S.M., Gruber, S., Lukas, S.E., Silveri, M.M., Hill, K.P., Killgore, W.D.S., Nickerson, L.D., 2020. Denoising scanner effects from multimodal MRI data using linked independent component analysis. Neuroimage 208.

Liu, S.Y., Thung, K.H., Qu, L.Q., Lin, W.L., Shen, D.G., Yap, P.T., 2021. Learning MRI artefact removal with unpaired data. Nature Machine Intelligence 3, 60–67.

Lusch, B., Kutz, J.N., Brunton, S.L., 2018. Deep learning for universal linear embeddings of nonlinear dynamics. Nature Communications 9.

Melzer, T.R., Keenan, R.J., Leeper, G.J., Kingston-Smith, S., Felton, S.A., Green, S.K., Henderson, K.J., Palmer, N.J., Shoorangiz, R., Almuqbel, M.M., Myall, D.J., 2020. Test-retest reliability and sample size estimates after MRI scanner relocation. Neuroimage 211, 116608.

Modanwal, G., Vellal, A., Buda, M., Mazurowski, M.A., 2020. MRI Image Harmonization using Cycle-Consistent Generative Adversarial Network. Medical Imaging 2020: Computer-Aided Diagnosis 11314.

Moyer, D., Steeg, G.V., Tax, C.M.W., Thompson, P.M., 2020. Scanner invariant representations for diffusion MRI harmonization. Magnetic Resonance in Medicine 84, 2174–2189.

Noble, S., Scheinost, D., Finn, E.S., Shen, X., Papademetris, X., McEwen, S.C., Bearden, C.E., Addington, J., Goodyear, B., Cadenhead, K.S., Mirzakhanian, H., Cornblatt, B.A., Olvet, D.M., Mathalon, D.H., McGlashan, T.H., Perkins, D.O., Belger, A., Seidman, L.J., Thermenos, H., Tsuang, M.T., van Erp, T.G.M., Walker, E.F., Hamann, S., Woods, S.W., Cannon, T.D., Constable, R.T., 2017a. Multisite reliability of MR-based functional connectivity. Neuroimage 146, 959–970.

Noble, S., Scheinost, D., Finn, E.S., Shen, X.L., Papademetris, X., McEwen, S.C., Bearden, C.E., Addington, J., Goodyear, B., Cadenhead, K.S., Mirzakhanian, H., Cornblatt, B.A., Olvet, D.M., Mathalon, D.H., McGlashan, T.H., Perkins, D.O., Belger, A., Seidman, L.J., Thermenos, H., Tsuang, M.T., van Erp, T.G.M., Walker, E.F., Hamann, S., Woods, S.W., Cannon, T.D., Constable, R.T., 2017b. Multisite reliability of MR-based functional connectivity. Neuroimage 146, 959–970.

Park, H.J., Friston, K., 2013. Structural and functional brain networks: from connections to cognition. Science 342, 1238411.

Poldrack, R.A., Gorgolewski, K.J., 2014. Making big data open: data sharing in neuroimaging. Nature Neuroscience 17, 1510–1517.

Pomponio, R., Erus, G., Habes, M., Doshi, J., Srinivasan, D., Mamourian, E., Bashyam, V., Nasrallah, I.M., Satterthwaite, T.D., Fan, Y., Launer, L.J., Masters, C.L., Maruff, P., Zhuo, C.J., Volzke, H., Johnson, S.C., Fripp, J., Koutsouleris, N., Wolf, D.H., Gur, R., Gur, R., Morris, J., Albert, M.S., Grabe, H.J., Resnick, S.M., Bryan, R.N., Wolk, D.A., Shinohara, R.T., Shou, H.C., Davatzikos, C., 2020. Harmonization of large MRI datasets for the analysis of brain imaging patterns throughout the lifespan. Neuroimage 208.

Radua, J., Vieta, E., Shinohara, R., Kochunov, P., Quide, Y., Green, M.J., Weickert, C.S., Weickert, T., Bruggemann, J., Kircher, T., Nenadic, I., Cairns, M.J., Seal, M., Schall, U., Henskens, F., Fullerton, J.M., Mowry, B., Pantelis, C., Lenroot, R., Cropley, V., Loughland, C., Scott, R., Wolf, D., Satterthwaite, T.D., Tan, Y., Sim, K., Piras, F., Spalletta, G., Banaj, N., Pomarol-Clotet, E., Solanes, A., Albajes-Eizagirre, A., Canales-Rodriguez, E.J., Sarro, S., Di Giorgio, A., Bertolino, A., Stablein, M., Oertel, V., Knochel, C., Borgwardt, S., du Plessis, S., Yun, J.Y., Kwon, J.S., Dannlowski, U., Hahn, T., Grotegerd, D., Alloza, C., Arango, C., Janssen, J., Diaz-Caneja, C., Jiang, W., Calhoun, V., Ehrlich, S., Yang, K., Cascella, N. G., Takayanagi, Y., Sawa, A., Tomyshev, A., Lebedeva, I., Kaleda, V., Kirschner, M., Hoschl, C., Tomecek, D., Skoch, A., van Amelsvoort, T., Bakker, G., James, A., Preda, A., Weideman, A., Stein, D.J., Howells, F., Uhlmann, A., Temmingh, H., Lopez-Jaramillo, C., Diaz-Zuluaga, A., Fortea, L., Martinez-Heras, E., Solana, E., Llufriu, S., Jahanshad, N., Thompson, P., Turner, J., van Erp, T., collaborators, E.C., 2020. Increased power by harmonizing structural MRI site differences with the ComBat batch adjustment method in ENIGMA. Neuroimage 218, 116956.

Rao, A., Monteiro, J.M., Mourao-Miranda, J., Initiative, A.D., 2017. Predictive modelling using neuroimaging data in the presence of confounds. Neuroimage 150, 23–49.

Schumann, G., Loth, E., Banaschewski, T., Barbot, A., Barker, G., Buchel, C., Conrod, P.J., Dalley, J.W., Flor, H., Gallinat, J., Garavan, H., Heinz, A., Itterman, B., Lathrop, M., Mallik, C., Mann, K., Martinot, J.L., Paus, T., Poline, J.B., Robbins, T.W., Rietschel, M., Reed, L., Smolka, M., Spanagel, R., Speiser, C., Stephens, D.N., Strohle, A., Struve, M., Consortium, I., 2010. The IMAGEN study: reinforcement-related behaviour in normal brain function and psychopathology. Molecular Psychiatry 15, 1128–1139.

Smallwood, R.F., Laird, A.R., Ramage, A.E., Parkinson, A.L., Lewis, J., Clauw, D.J., Williams, D.A., Schmidt-Wilcke, T., Farrell, M.J., Eickhoff, S.B., Robin, D.A., 2013. Structural brain anomalies and chronic pain: a quantitative meta-analysis of gray matter volume. J Pain 14, 663–675.

Tanaka, S.C., Yamashita, A., Yahata, N., Itahashi, T., Lisi, G., Yamada, T., Ichikawa, N., Takamura, M., Yoshihara, Y., Kunimatsu, A., Okada, N., Hashimoto, R., Okada, G., Sakai, Y., Morimoto, J., Narumoto, J., Shimada, Y., Mano, H., Yoshida, W., Seymour, B., Shimizu, T., Hosomi, K., Saitoh, Y., Kasai, K., Kato, N., Takahashi, H., Okamoto, Y., Yamashita, O., Kawato, M., Imamizu, H., 2021. A multi-site, multi-disorder resting-state magnetic resonance image database. Scientific Data 8.

Tong, Q.Q., Gong, T., He, H.J., Wang, Z., Yu, W.W., Zhang, J.J., Zhai, L.H., Cui, H.S., Meng, X., Tax, C.W.M., Zhong, J. H., 2020. A deep learning-based method for improving reliability of multicenter diffusion kurtosis imaging with varied acquisition protocols. Magnetic Resonance Imaging 73, 31–44.

Tong, Q.Q., He, H.J., Gong, T., Li, C., Liang, P.P., Qian, T.Y., Sun, Y., Ding, Q.P., Lie, K.C., Zhong, J.H., 2019. Reproducibility of multi-shell diffusion tractography on traveling subjects: A multicenter study prospective. Magnetic Resonance Imaging 59, 1–9.

van den Heuvel, M.P., Sporns, O., 2019. A cross-disorder connectome landscape of brain dysconnectivity. Nat Rev Neurosci 20, 435–446.

Vovk, U., Pernus, F., Likar, B., 2007. A review of methods for correction of intensity inhomogeneity in MRI. Ieee Transactions on Medical Imaging 26, 405–421.

Xia, M., He, Y., 2017. Functional connectomics from a “big data” perspective. Neuroimage 160, 152–167.

Xia, M.R., Si, T.M., Sun, X.Y., Ma, Q., Liu, B.S., Wang, L., Meng, J., Chang, M., Huang, X.Q., Cheni, Z.Q., Tang, Y.Q., Xu, K., Gong, Q.Y., Wang, F., Qiu, J., Xie, P., Li, L.J., He, Y., Wor, D.M.D.D., 2019. Reproducibility of functional brain alterations in major depressive disorder: Evidence from a multisite resting-state functional MRI study with 1,434 individuals. Neuroimage 189, 700–714.

Yamashita, A., Yahata, N., Itahashi, T., Lisi, G., Yamada, T., Ichikawa, N., Takamura, M., Yoshihara, Y., Kunimatsu, A., Okada, N., Yamagata, H., Matsuo, K., Hashimoto, R., Okada, G., Sakai, Y., Morimoto, J., Narumoto, J., Shimada, Y., Kasai, K., Kato, N., Takahashi, H., Okamoto, Y., Tanaka, S.C., Kawato, M., Yamashita, O., Imamizu, H., 2019. Harmonization of resting-state functional MRI data across multiple imaging sites via the separation of site differences into sampling bias and measurement bias. Plos Biology 17.

Yu, M.C., Linn, K.A., Cook, P.A., Phillips, M.L., McInnis, M., Fava, M., Trivedi, M.H., Weissman, M.M., Shinohara, R.T., Sheline, Y.I., 2018. Statistical harmonization corrects site effects in functional connectivity measurements from multi-site fMRI data. Human Brain Mapping 39, 4213–4227.

Zhao, F.Q., Wu, Z.W., Wang, L., Lin, W.L., Xia, S.R., Shen, D.G., Li, G., C, U.U.B.C.P., 2019. Harmonization of Infant Cortical Thickness Using Surface-to-Surface Cycle-Consistent Adversarial Networks. Medical Image Computing and Computer Assisted Intervention - Miccai 2019, Pt Iv 11767, 475–483.

Zuo, L.R., Dewey, B.E., Liu, Y.H., He, Y.F., Newsome, S.D., Mowry, E.M., Resnick, S.M., Prince, J.L., Carass, A., 2021. Unsupervised MR harmonization by learning disentangled representations using information bottleneck theory. Neuroimage 243.

