## Supplementary material for "A deep learning-based multisite neuroimage harmonization framework established with traveling-subject dataset": SI Appendix

**Supplementary Information**

### Competing methods

To verify the effectiveness of site effect adjustment, we compared DeRed with other harmonization methods including general linear model (GLM) harmonization, global scaling (GS) harmonization, and ComBat harmonization.

### GLM method

The GLM harmonization method assesses site differences by fitting a linear regression model that regards site labels as unique, independent variables. The linear regression model can be written as:

$$y_{ijv} = const + X_s^T s^{GLM} + \varepsilon_{ijv} \tag{5.1}$$

where $y_{ijv}$ denotes the GMV measurement of the *i-th* site, *j-th* subject, and *v-th* voxel. *const* is an average measurement over all voxels. $X_s$ represents a dummy encoding, $s^{GLM}$ represents a n×1 vector of the estimated differences of n sites, and $\varepsilon_{ijv}$ denotes a residual term. The GM volume measurements were adjusted by removing estimated differences of n sites. In summary, the harmonized GM volume measurements were formulated as follows:

$$y_{ijv}^{GLM} = y_{ijv} - X_s^T \hat{s}^{GLM} \tag{5.2}$$

where $\hat{s}^{GLM}$ denotes estimated differences of n sites.

### GS method

The GS harmonization method assesses the discrepancy between the average GM volume measurement of subjects from a single site and that of subjects from all sites. The discrepancy representation can be expressed by additive and multiplicative parameters, which can be estimated by fitting a linear model:

$$\bar{y}_{i,v} = \theta_{i,location} + \theta_{i,scale}\bar{y}_v + \varepsilon_{ijv} \tag{6.1}$$

where $\theta_{i,location}$ and $\theta_{i,scale}$ represent additive and multiplicative parameters for the *i-th* site, respectively, $\bar{y}_{i,v}$ denotes the average GM volume measurement of the *v-th* voxel for site *i* and $\bar{y}_v$ denotes the average GM volume measurement of the *v-th* voxel for all sites. After performing ordinary least squares (OLS), the estimated parameters $\hat{\theta}_{i,location}$ and $\hat{\theta}_{i,scale}$ can be used to adjust the site effect as follows:

$$y_{ijv}^{GS} = \frac{y_{ijv} - \hat{\theta}_{i,location}}{\hat{\theta}_{i,scale}} \tag{6.2}$$

### ComBat method

The ComBat harmonization method is currently widely used in multisite data correction and has shown excellent results in the harmonization task for cortical thickness measurements across scanners and sites. The ComBat approach considers that the data differences caused by biological factors can be estimated by the biological subject information, such as age and sex. if the GM volume data still differ between sites after regression of biological factors in the linear models, these differences are influenced by site-related effects. These differences were adjusted by applying empirical Bayesian estimation which can improve model effectiveness for small sample sizes. The ComBat model can be written as:

$$y_{ijv} = \alpha_v + X_{ij}^T \beta_v + \gamma_{iv} + \delta_{iv}\varepsilon_{ijv} \tag{7.1}$$

where $\gamma_{iv}$ represents an additive parameter for the *i-th* site and *v-th voxel*, $\delta_{iv}$ represents a multiplicative parameter for the *i-th* site and *v-th* voxel, and $X_{ij}$ refers to a design matrix involving to covariates of interest; and only age was considered in this work. $\alpha_v$ is the averaged GM volume measurement of the *v-th* voxel, and $\beta_v$ is a vector of the regression coefficients. The harmonized GM volume measurements are given by:

$$y_{ijv}^{ComBat} = \frac{y_{ijv} - \hat{\alpha}_v - X_{ij}^T\hat{\beta}_v - \hat{\gamma}_{iv}}{\hat{\delta}_{iv}} + \hat{\alpha}_v + X_{ij}^T\hat{\beta}_v \quad (7.2)$$

in which $\hat{\alpha}_v$, $\hat{\beta}_v$, $\hat{\gamma}_{iv}$ and $\hat{\delta}_{iv}$ are represent estimated parameters.

### Figure S1

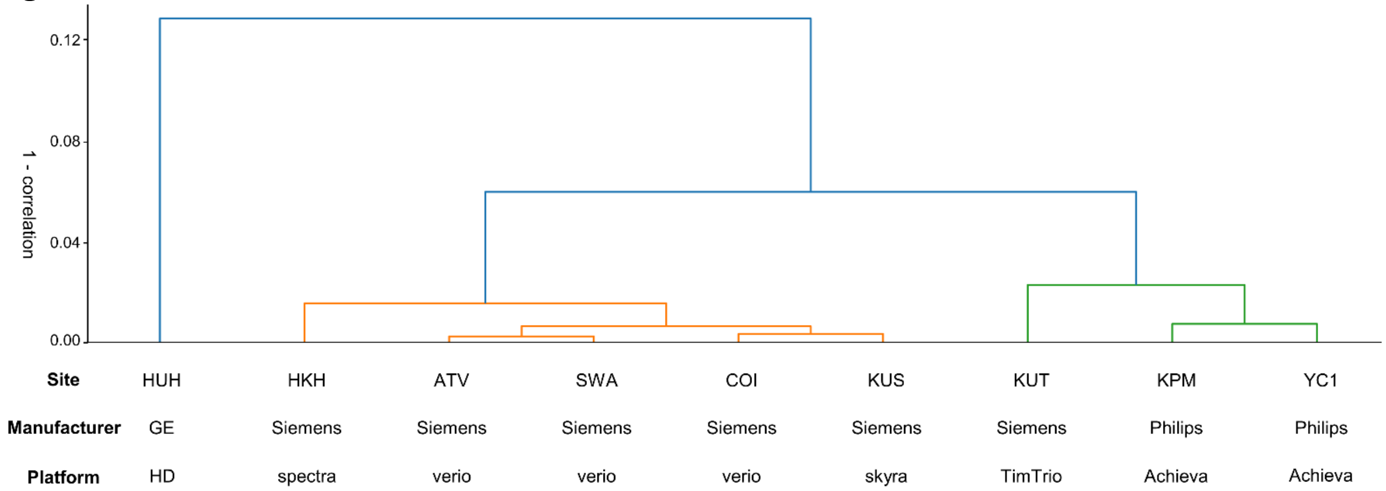


**Fig. S1 Hierarchical clustering dendrogram for averaged GMV map.** The height of each linkage indicates the difference calculated by (1 − correlation) between the clusters joined by that link. The site effects are mainly divided into clusters according to an order of MRI manufacturer and scanner type.

### Figure S2

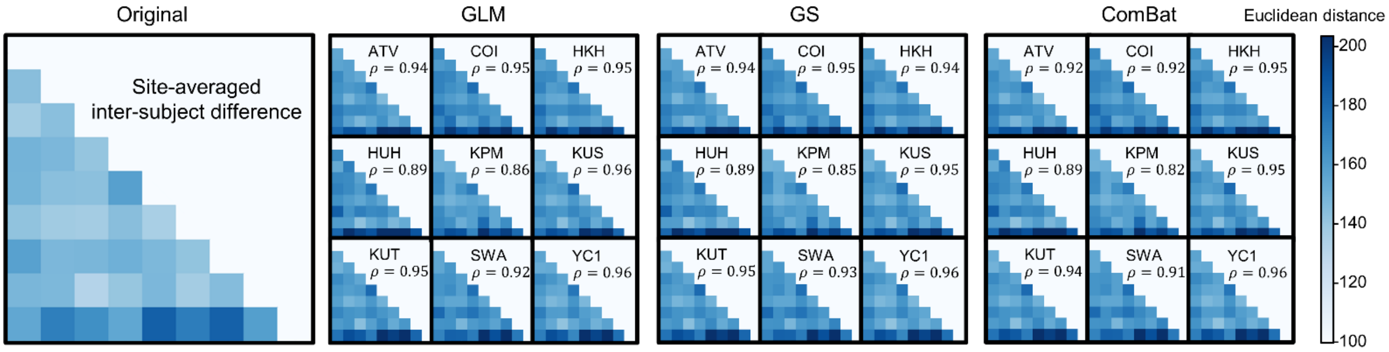


**Fig. S2 Inter-subject difference matrix after different harmonization methods.** The difference matrices were averaged across sites for original data. The color depth of *i-th* row and *j-th* column grid for each matrix represents the Euclidean distance between the *i-th* and *j-th* subjects. Spearman's correlation coefficients are illustrated by $\rho$ ($p$<0.001).
